# tRNA modifications inform tissue specific mRNA translation and codon optimization

**DOI:** 10.1101/2023.10.24.563884

**Authors:** Daisuke Ando, Sherif Rashad, Thomas J Begley, Hidenori Endo, Masashi Aoki, Peter C Dedon, Kuniyasu Niizuma

**Affiliations:** Department of Neurology, Tohoku University Graduate School of Medicine, Sendai, Japan; Department of Neurosurgical Engineering and Translational Neuroscience, Tohoku University Graduate School of Medicine, Sendai, Japan; Department of Neurosurgical Engineering and Translational Neuroscience, Graduate School of Biomedical Engineering, Tohoku University, Sendai, Japan; Department of Biological Sciences, University at Albany, Albany, NY, USA; Department of Neurosurgery, Tohoku University Graduate School of Medicine, Sendai, Japan; Department of Biological Engineering, Massachusetts Institute of Technology, MA, USA

**Keywords:** tRNA modifications, Queuosine, Codon optimality, Ribosome profiling, Codon optimization, mRNA translation

## Abstract

The tRNA epitranscriptome has been recognized as an important player in mRNA translation regulation. Our knowledge of the role of tRNA epitranscriptome in fine-tuning translation codon decoding at tissue or cell levels remains incomplete. Here, we analyzed seven tissues from mice for the expression of tRNA modifications and mature tRNAs as well as mRNA translation and codon decoding. Our analysis revealed distinct enrichment patterns of tRNA modifications in tissues. Queuosine (Q) tRNA modification was most enriched in the brain compared to other tissues, while mitochondrial tRNA modifications and tRNA expression was highest in the heart. Using three different metrics for codon analysis; isoacceptors frequencies, total codon frequencies, and A-site pausing, we revealed a strong bias towards A/T ending codons in most tissues except for the brain. Using this observation, we synthesized, and delivered *in vivo*, codon mutated EGFP for Q-codons, where the C-ending Q-codons were replaced with U-ending codons. The protein levels of mutant EGFP were downregulated in liver, which is poor in Q, when NAC codons were exchanged for NAU codons, while in brain EGFP levels did not change. This data shows that understanding tRNA modifications enrichments across tissues is not only essential for understanding codon decoding and bias, but it can also be utilized for optimizing gene and mRNA therapeutics to be more tissue, cell, or condition specific.

## Introduction

The translation of the gene code to functional protein is at the heart of all processes of life. This multistep process that starts from transcription all the way to post-translational protein modifications is under strict regulation at every step. At the heart of this process lies mRNA translation, which is one of the most complex and heavily regulated processes. mRNA modifications and structure(1, 2), ribosome assembly and modifications(3), and tRNA epitranscriptome(4) are few of the factors that interplay to regulate proper mRNA translation and proteostasis. In recent years, there has been an increased interest in the complexity and regulatory roles of tRNA epitranscriptome on mRNA translation(4, 5). Importantly, alteration of tRNA expression(6, 7), tRNA derived small non-coding RNAs (tsRNAs)(8), and tRNA modifications(9) were shown to impact mRNA translation and play roles in diseases such as cancer, metabolic disorders, and neuropsychiatric disorders(7–12). However, our understanding of how tRNA epitranscriptome regulates mRNA translation, while substantially improved over the past years, is far from complete.

A limited set of tRNA anticodons, 48 in humans and 47 in mice, decode 64 codons in the mRNA(13). To compensate for this mismatch, non-cognate codon recognition at the wobble anti-codon position, position 34 in the tRNA, allows the decoding of the codons via the limited available tRNA pool(4, 14). This process is fine-tuned by tRNA modifications at the wobble position(4, 5, 15). For example, tRNA modifications can expand or restrict the codon decoding by anticodons(4). A famous example is Adenosine-to-Inosine conversion (A-to-I) which expands the recognition of NNU codons by ANN anticodons to NNU, NNC, and NNA by INN anticodons(4, 16) or Queuosine modifications that expands the recognition of NAC codons to NAY (NAC and NAU) codons(17–19).

Previous works have elucidated the dynamic nature of tRNA epitranscriptome in regulating processes such as the stress response(6, 15, 20–23) and growth conditions(24) and how these dynamic changes impact codon recognition(6, 23, 25). Nonetheless, much remains to be elucidated on the role of tRNA epitranscriptome in regulating physiologic mRNA translation in various tissues and cells, and their role in tissue physiology. Several groups have analyzed tRNA modifications(26) and tRNA expression(27, 28) in mice tissues in order to elucidate such regulation. While these efforts did reveal important phenomena on tissue specific expression of tRNA epitranscriptome and its potential links to mRNA translation, they were limited in focusing on only one aspect of tRNA epitranscriptome at a time or not providing direct correlation with mRNA translation and codon decoding. In addition, the codon metrics used for the analysis in some works still require revisiting.

Here, we analyzed tRNA epitranscriptome (tRNA modifications and mature tRNA expression) from 7 mouse tissues using sequencing and mass spectrometry [Figure 1a]. We combined our analysis with Ribo-seq analysis and applied various codon metrics to our dataset to reveal that tRNA modifications are linked to codon usage and optimality in various tissues. We further utilized the expression of tRNA modifications as an input for codon optimization of gene constructs to show how such information can be used to fine-tune mRNA delivery and viral-based gene therapeutics in a tissue and cell specific manner.

**Figure 1:**
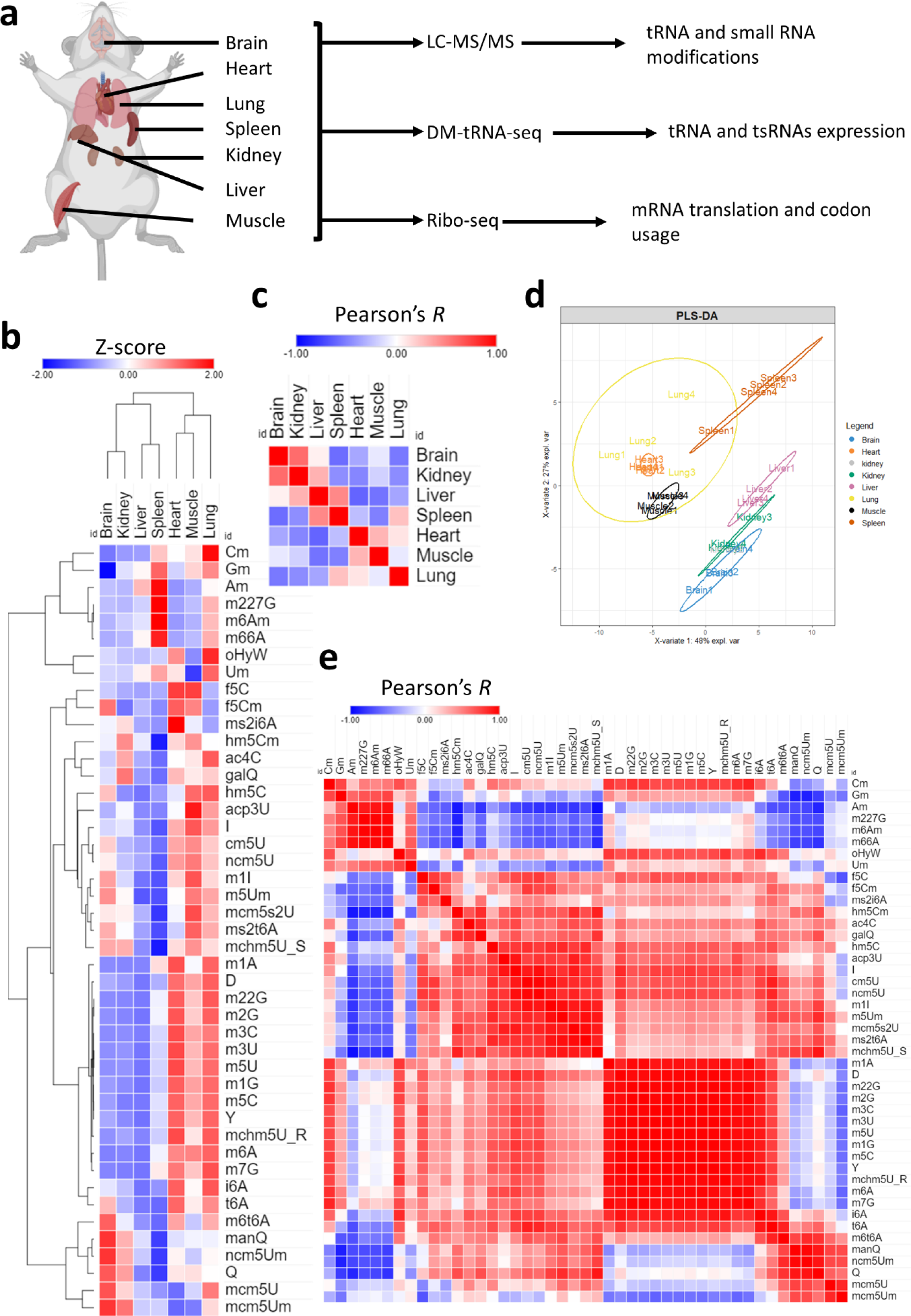
Analysis of tRNA modifications landscape across tissues. a: Schematic showing the tissues analyzed and methods used in this study. b: Expression of 45 different modifications across the 7 tissues. Data presented as Z-scores across tissues. N = 4 biological replicates per group. c: Pearson’s correlation between different tissues using z-scores of tRNA modifications as input. d: PLS-DA analysis of tissue clustering patterns using normalized peak areas of tRNA modifications as input. e: Pearson’s correlation across modifications.

## Results

### ⮚ Tissue specific tRNA modification enrichment

We first began by exploring the tRNA modifications landscape in each of the 7 tissues used in this study [Figure 1a]. Using high throughput LC-MS/MS analysis of tRNA enriched small RNA fraction (>90% tRNA), we were able to detect 45 known mammalian modifications that occur in the tRNA or in other small non-coding RNAs (such as m6A and m66A(4, 29)) [Supplementary figure 1a](4). Our results revealed significant tissue specific enrichment of certain modifications as well as tissue and modification clustering patterns [Figure 1b, supplementary table 1]. A cluster of related modifications, mcm5U, mcm5Um, and ncm5Um, which are regulated by the same enzyme system(4, 30) were most enriched in the brain as well as queuosine (tRNA-Q) and its derivative mannosyl queuosine (manQ) but not its galactosyl derivative (galQ). We also observed that f5C and f5Cm, which exist in mitochondrial tRNA-Met and cytosolic tRNA-Leu^CAA^ (^31^) were most enriched in the heart and muscle. Another mitochondrial tRNA specific modification, ms2i6A, was most enriched in the heart. On the other hand, the liver and spleen had lower levels of almost all modifications, albeit the spleen showed an enrichment in a specific set of modifications that include 2’O-ribose modified nucleotides such as Cm, Gm, and Am, as well as several modifications not known to occur in the tRNA such as m66A and m6Am(4, 29) but occur in other small non-coding RNAs.

Next, we directed our attention to how tissues cluster and correlate with each other, which can give an idea about the fine-tuning of tRNA modifications towards tissue specific functions. To that end, we conducted Pearson’s correlation [Figure 1c] and partial least square regression discriminant analysis (PLS-DA) [Figure 1d] to understand such relations between tissues. Peason’s correlation revealed that the brain had higher correlation with the kidney while it had a negative correlation with the spleen and lungs and no correlation with the liver [Figure 1c]. PLS-DA analysis [Figure 1d] confirmed the relationship between tissues, revealing that the brain clusters closely with the kidney while the lung, spleen, muscles, and heart formed another cluster. Further analysis of the distribution of modifications in the PLS-DA plot [Supplementary figure 1b], revealed how clusters of modifications correlated with tissue clusters. In addition, VIP analysis [Supplementary figure 1c] revealed important contributors to each tissue cluster expression pattern. For example, m66A contributed to the Spleen and Lung clustering patterns while tRNA-Q contributed to the brain followed by kidneys. In addition to tissue clustering, we analyzed the correlation between tRNA modifications [Figure 1e]. Our analysis revealed clusters of modifications that follow the same tissue expression patterns. For example, methylation modifications, such as m1A, m5C, m3C, etc., generally clustered together, while 2’O-ribose modifications clustered together and so on.

In summary, our analysis revealed significant tissue specific enrichment patterns of tRNA modifications.

### ⮚ Mature tRNAs and anticodons are expressed equally across tissues

Here, we analyzed the expression of mature tRNAs and their respective anticodons across tissues. Sequencing analysis revealed tissue specific variations in the expression of tRNAs and various non-coding RNAs. For example, spleen and lung showed very low levels of tRNAs compared to other tissues, while they had significantly higher snRNAs and snoRNAs [Figure 2a]. This pattern was also evident when RNA samples were analyzed using Bioanalyzer. Spleen and lung consistently showed higher fractions of small RNA peaks between 20∼40 nucleotides in length [Supplementary figure 2]. Such patterns cannot be attributed to simple RNA degradation during extraction as they were operator independent (replicated by 2 different operators) and were found in all samples analyzed. This finding can also explain the enrichment of non-tRNA modifications in the spleen and lung compared to other tissues. In addition, mitochondrial tRNAs (Mt_tRNA) were expressed in the heart at much higher levels, consistent with the enrichment of mitochondria related tRNA modifications in the heart. We also observed notable differences in the ratios of tRNAs pertaining to specific amino acids in different tissues. Notably, Selenocysteine (SeC) representation was higher in spleen and lung, which might be related to the known immune functions of SeC (32). Next, we collapsed the mature tRNA counts by anticodon and calculated the expression of different tRNAs. As reported previously(27), the relative normalized expression of different tRNA isodecoders within each tissue were relatively stable, which is reflective of the stability of tRNA housekeeping genes(27, 33) [Figure 2c]. However, when comparing the relative expression of each isodecoder across different tissues [Figure 2d] differences start to appear. Indeed, given the low expression of tRNAs in the spleen, almost all isodecoders were expressed at lower levels compared to other tissues, except for Ser^CGA^. Further, despite the representation of SeC amino acid related counts in the spleen and lungs [Figure 2b], the isodecoder for SeC (SeC^TCA^) was most expressed in the kidney. We also analyzed the expression of tRNA-Q modified tRNAs. Asp^GTC^ was most enriched in the kidneys, while Asn^GTT^ was equally enriched in both the Kidney and brain. On the other hand, His^GTG^ was equally expressed across tissues except in lung and spleen where it was expressed at lower levels. Paradoxically, Tyr^GTA^ was most expressed in the liver and underrepresented in the brain. This can be explained, however, by the lower expressed of galactosyl-queuosine modification in the brain which occurs in Tyr^GTA^ (4, 19). Pearson’s correlation analysis using data from Figure 2d [Figure 2e] revealed a different pattern of clustering compared to what was observed at the level of tRNA modifications. For example, the brain had modest positive correlation with the heart and liver, while it correlated negatively with the kidneys. These changes, however, remain subtle, as when analyzing the correlation between tissues using normalized read counts (data from figure 2c), there is near perfect correlation [Figure 2f].

**Figure 2:**
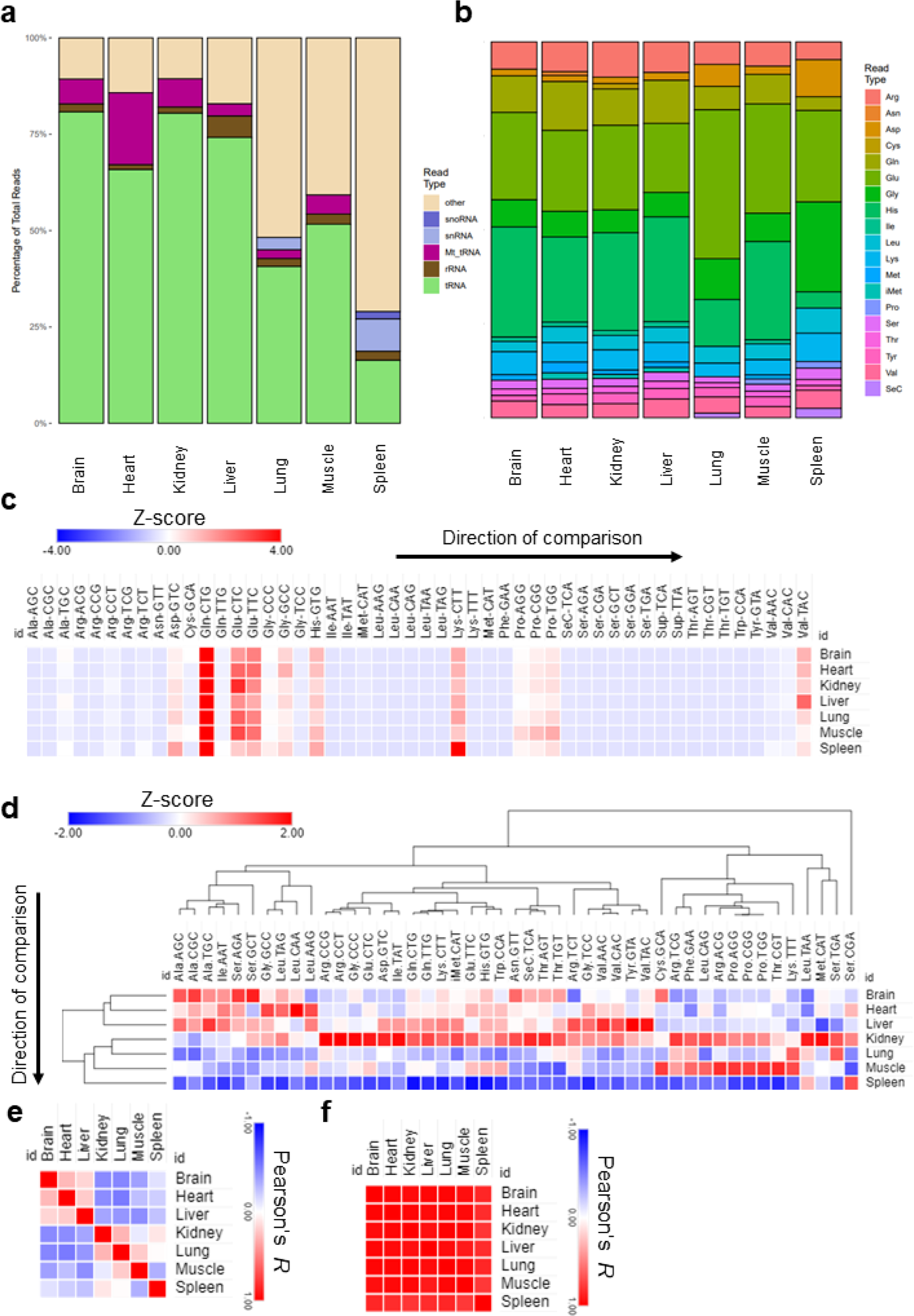
Analysis of mature tRNA expression across tissues. a: Distribution of detected small non-coding RNAs in the sequencing datasets. b: Distribution of different amino acids representing tRNAs in the datasets. c: Expression of different mature tRNAs in our dataset normalized by their normalized read counts and represented as Z-score. Here, z-scores were calculated for each tissue separately. d: Z-scores of mature tRNA expression across different tissues. e: Pearson’s correlation coefficient between tissues based on Figure 2d. f: Pearson’s correlation coefficient analysis between tissues based on figure 2c. N = 3 biological replicates per group.

While the isoacceptors cytosolic tRNAs normalized expression were nearly uniform across tissues, mitochondrial tRNAs revealed a difference picture. The heart showed much higher normalized read counts compared to all other tissues for almost all detected Mt-tRNAs with only a few exceptions [Supplementary figures 3a and 3b] and tissue specific clustering was observed [Supplementary figure 3c].

In summary, our analysis, in agreement with previous work(27), revealed almost uniform normalized expression of tRNA isodecoders in different tissues, in agreement with the notion of the existence of tRNA housekeeping genes that are stably expressed in all cells and tissues(33).

### ⮚ Unique translational patterns observed in various tissues

To understand the potential links between the tRNA epitranscriptome and tissues specific translational patterns and codon decoding, we conducted Ribosome profiling (Ribo-seq). We were mainly interested in observing the clustering patterns at the translational level as well as conducting codon analytics to corroborate these data with our observations at the tRNA level. We used various ways to cluster tissues at the translational level; heatmap clustering [Figure 3a, supplementary figure 4a], tSNE [Figure 3b], and Gene level UMAP [Figure 3c], all revealed unique clustering of the brain vs other tissues. It was also evident that the clustering patterns observed at the translational level could faithfully replicate those observed at the tRNA modifications level [Figures 1c and 1d], where the brain was somewhat unique but closer to kidneys and liver, while the spleen and lung clustered closely together as well as the heart and muscles. We next analyzed the differentially translated genes (DTGs) in each tissue versus all the others [Figure 3d] and observed various gene translational patterns. The brain showed the largest number of significant DTGs versus other tissues followed by the liver and heart. Indeed, this was also evident when we conducted detailed DTG analysis for tissue pairs [Supplementary table 2]. Using the DTGS from each tissue versus all others, we generated a UMAP of gene ontology biological processes (GOBP) pathway analysis, which showed using clustering for each tissue in terms of enriched pathways, with significant overlaps between closely related tissues (Heart and muscle or lung and spleen) [Supplementary figure 4b-c]. In addition, the activation matrix of GOBP pathways showed unique pathway enrichment that also followed the clustering patterns of tissues [Figure 3e]. For example, pathways related to mitochondrial function and muscle contraction were enriched in the heart and muscles, in agreement with the enrichment of tRNA modifications and Mt-tRNAs expression. Finally, we evaluated whether the clustering and gene translation patterns in different tissues can be used to accurately annotate said tissues. The analysis revealed a good degree of success in annotating the tissues, further supporting the robustness of the sequencing dataset [Supplementary figure 5]. We further explored our Ribo-seq data for the expression of known tRNA and small RNA modifying enzymes, with a focus on tRNA-Q and mcm5U related enzymes(4) [Supplementary figure 6a]. While we observed differences in expression across tissues, clustering, enzyme expression, as well as PLS-DA analysis did not correlate with what we observed at the level of tRNA modifications themselves [Supplementary figure 6b], indicating that the changes observed at the modifications’ levels are not mainly driven by differences in enzyme expression between tissues.

**Figure 3:**
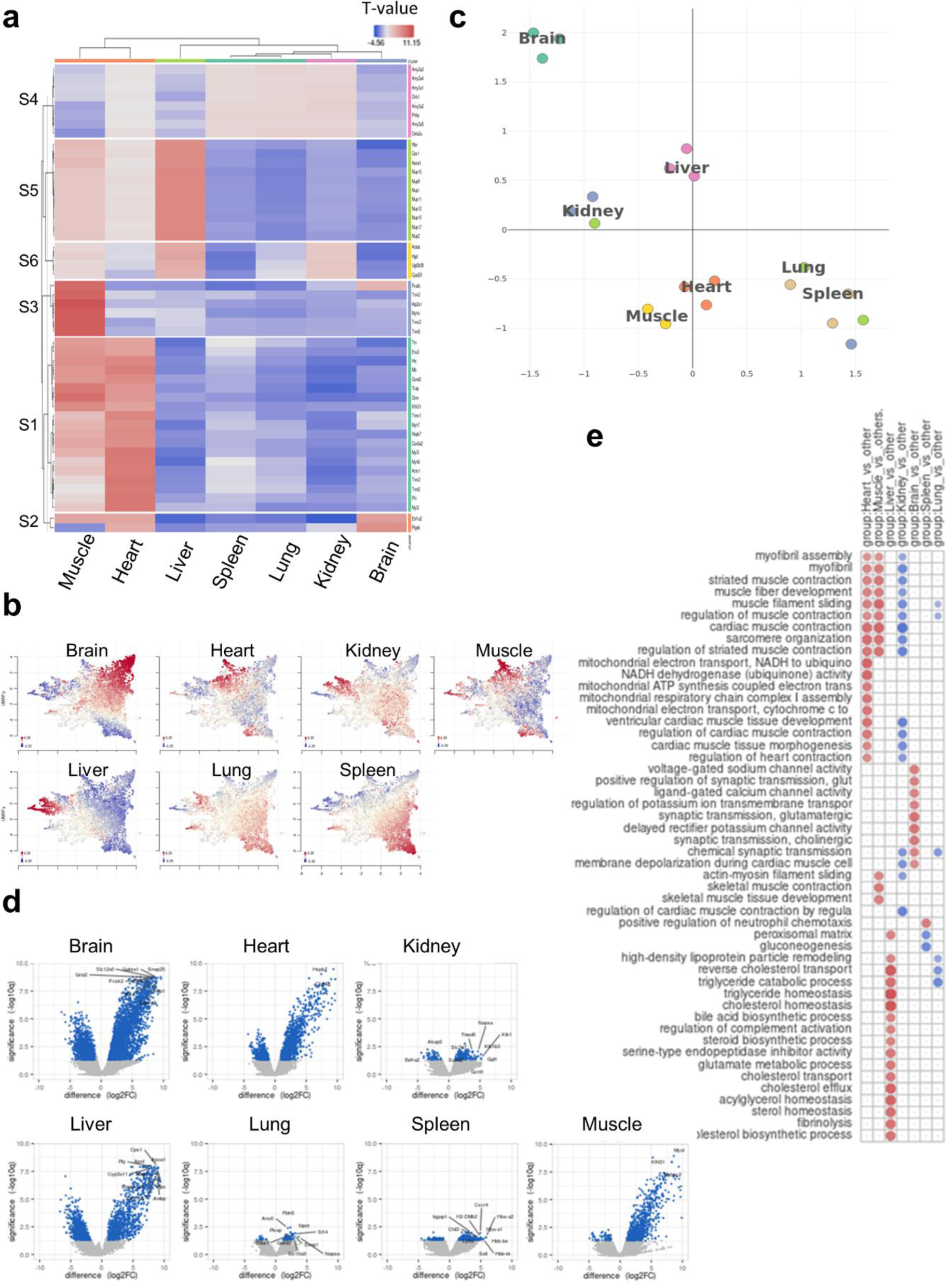
Ribo-seq analysis of tissue specific translational patterns. a: Heatmap clustering between tissues (Cluster annotation is in Supplementary figure 4a). b: Gene-level UMAP analysis. c: tSNE clustering analysis. d: Differentially translated genes (DTGs) analysis for each tissue versus all other tissues. e: GOBP activation matrix of pathway enrichment using input from **d**. N = 3 biological replicates per group.

### ⮚ Tissue codon usage and optimality are governed by tRNA modifications levels

tRNA modifications are known to drive mRNA translation by altering codon recognition, usage, and optimality(4, 17, 23, 25). To elucidate the potential for such regulatory fine-tuning on tissue specific mRNA translation, we analyzed three metrics for codon analysis; isoacceptors codon frequency (which measures the enrichment of synonymous codons compared to each other), total codon frequency (which compares the expression of a codon with all other codons in the mRNA sequence)(34, 35), and A-site pausing (which measures ribosome dwelling at each codon)(36) [Supplementary figure 7].

First, we started by analyzing the codon isoacceptors frequencies in each tissue using the top and bottom 200 expressed genes by normalized read counts [Figure 4]. Heatmap clustering revealed a distinctive A/T vs G/C ending codon bias with few exceptions [Figure 6a, supplementary figure 8a]. Except for the brain and the heart, all tissues were biased towards A/T-ending codons, evident from the increased usage of these codons in upregulated genes and their underusage in downregulated genes. On the other hand, the brain and heart showed the least degree of bias and showed a shift towards G/C-ending codons selection. Pearson’s correlation coefficient analysis was done on the isoacceptors frequencies of the top and bottom 200 genes separately [Figure 4b]. It was evident that the differences in tissues were mainly apparent in the downregulated genes, with the brain being furthest from all tissues except the heart, in agreement with the observed codon biases. Further, PLS-DA analysis revealed unique clustering patterns akin to what was observed at the level of tRNA modifications and translation, with few exceptions. For example, the spleen and lung clustered together and were far from the brain [Figure 4c, supplementary figure 8b]. Based on the impression from our data that the brain is unique compared to other tissues, we focused our attention on 2 tRNA modifications clusters which were highly expressed in the brain; tRNA-Q and its derivatives mannosyl-queuosine (manQ) and galactosyl-queuosine (gal-Q) and the mcm5U cluster and its related modifications and their corresponding codons. First, we explored the usage of Q-decoded codons and their correlation with the expression of tRNA-Q [Figure 4d]. tRNA-Q expands the codon decoding from NAC to NAY codons(15), which includes Asn-AAC/U, Asp-GAC/U, His-CAC/U, and Tyr-TAC/U. manQ decodes Asp-GAC/U while galQ decodes Tyr-TAC/U. We observed a positive correlation between C-ending Q-codons and Q modifications, except for Tyr-TAC. This can be explained by the fact that Q and manQ were enriched in the brain but galQ did not show such enrichment [Figure 1b]. In addition, Tyr-TAC/U did not show a strong bias in our analysis as compared to other Q-codons [Figure 4a]. It is also notable to mention that the C-ending Q-codons, NAC codons, were negatively correlated with their synonymous NAU codons, signifying strong codon bias and selection [Figure 4e]. On the other hand, for mcm5U and mcm5s2U modifications, which are related to NAA codons recognition, mcm5U showed a positive correlation trend with GlnCAA and LysAAA, while mcm5s2U showed a negative correlation trend with LysAAA [Figure 4f], and the correlation coefficient for the other NAA codons was generally low, compared to Q codon.

**Figure 4:**
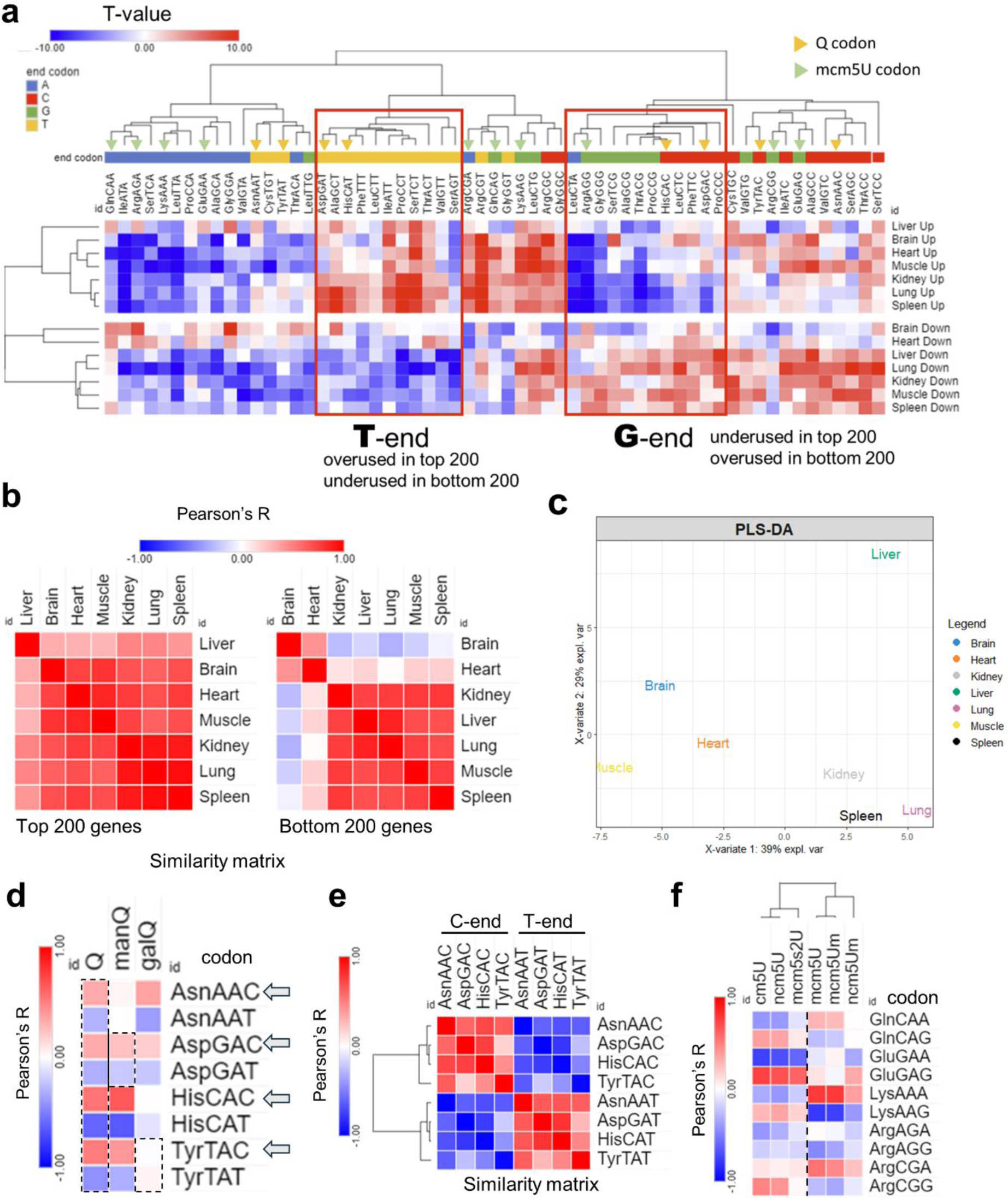
Analysis of isoacceptors codon usage. a: Heatmap of isoacceptors codon frequency in each tissue. Isoacceptors frequencies T-stats were calculated using top 200 (up) or bottom 200 (down) expressed genes from the Ribo-seq data in each tissue. Data presented as T-stat versus the genome average. T-stat of >2 or <-2 indicates *p <* 0.05. b: Heatmap of Pearson’s correlation coefficients among 7 tissues by codon frequency from top (left) and bottom (right) 200 genes. c: PLS-DA analysis of tissue clustering patterns using the top 200 genes. d: Pearson’s correlation analysis between codon frequencies (T-stat of isoacceptors frequencies) and Q modification enrichment (z-score of tRNA-Q across tissues). e: Pairwise correlation analysis of Q codon from figure 4a. g, f: Pearson’s correlation analysis of codon frequency and mcm5U and related modification enrichment.

Next, we repeated our analysis at the level of total codon usage, which can give us clues as to the mRNA amino acids and sequence preferences [Figure 7]. In this analysis, we observed strong bias in all tissues between codon frequencies [Figure 5a], but with weaker A/T vs G/C ending codon bias. Differences between tissues were apparent in the bottom 200 genes and not the top 200 genes when we conducted Pearson’s correlation coefficient analysis [Figure 5b]. In addition, PLS-DA analysis, while the tissues showed different clustering patterns compared to isoacceptors frequencies, still showed the brain to be unique while the lung and spleen clustered together [Figure 5c and supplementary figure 9]. The correlation between Q-codons and mcm5U-codons with their respective modifications was weaker or disappeared altogether, and there was no bias observed between isoacceptors in their total codon frequencies counts [Figure 5d-f].

**Figure 5:**
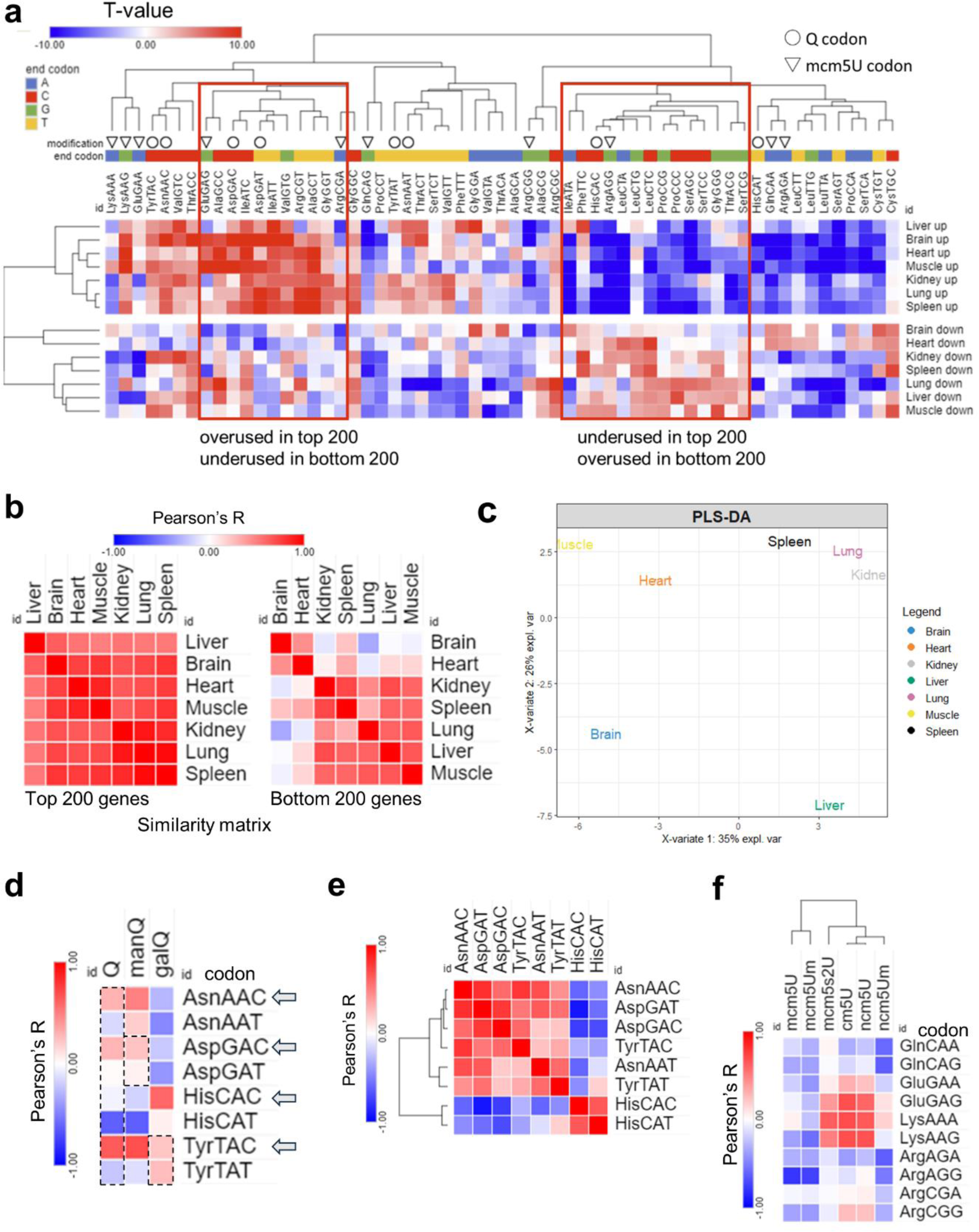
Analysis of total codon frequency. a: Heatmap of total codon frequency in each tissue. T-stats were calculated using frequency of total codon counts using the same input as 4a. b: Heatmap of Pearson’s correlation coefficients among 7 tissues by codon frequency from top (left) and bottom (right) 200 genes. c: PLS-DA analysis of tissue clustering patterns using the top 200 genes. d: Pearson’s correlation analysis between codon frequencies (T-stat of isoacceptors frequencies) and Q modification enrichment (z-score of tRNA-Q across tissues). e: Pairwise correlation analysis of Q codon from figure 5a. g, f: Pearson’s correlation analysis of codon frequency and mcm5U and related modification enrichment.

**Figure 6:**
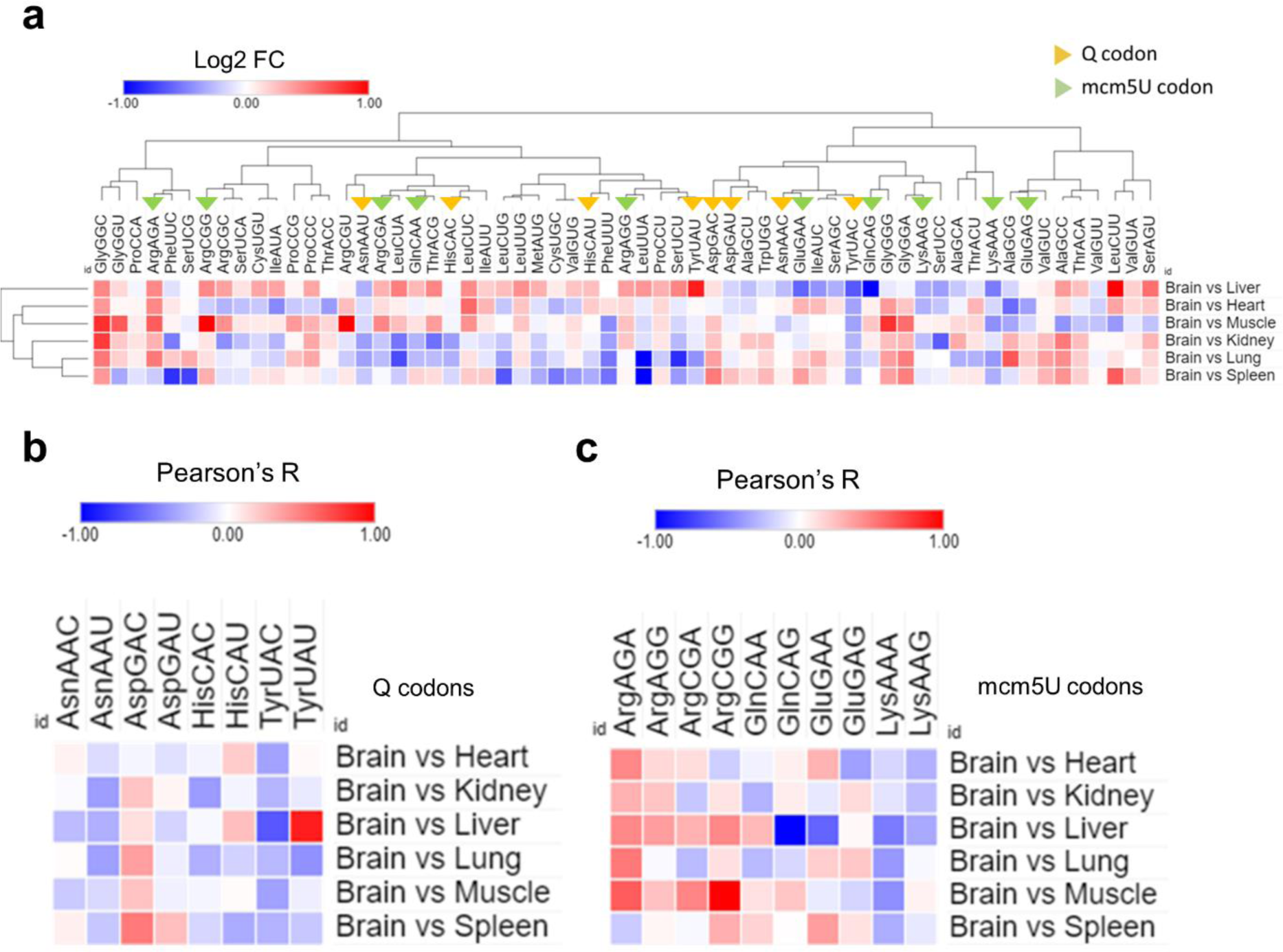
Analysis of A-site ribosome pausing. a: Heatmap of A-site ribosome pausing analysis. The relative dwelling time in the brain compared to other tissues is represented as log 2-fold change (log2FC). b, c: Selection of the codons related to Q modifications (c) or mcm5U modifications (d) from figure 8a for clarity.

**Figure 7:**
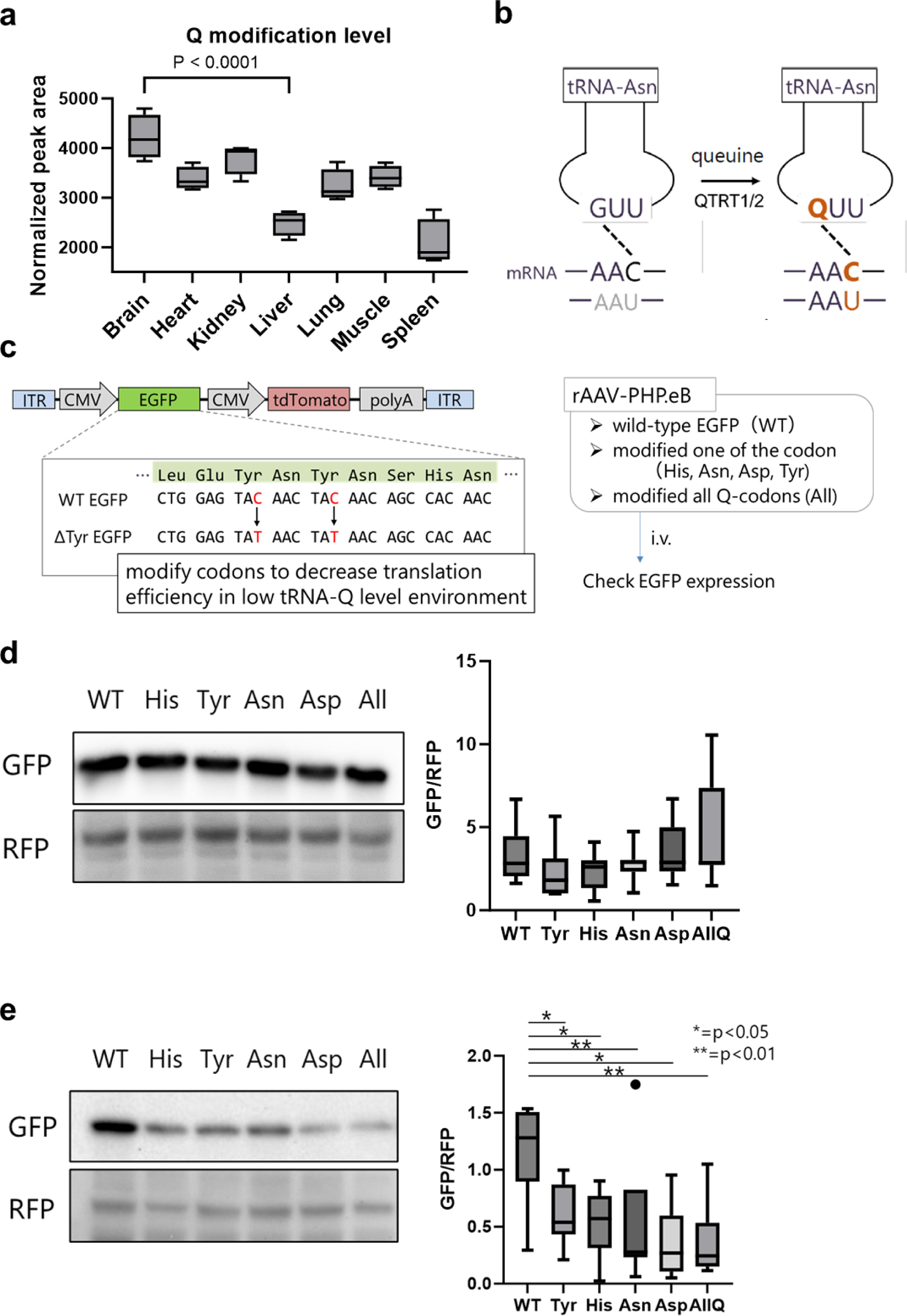
tRNA modifications inform codon optimization algorithms. a: Expression of tRNA-Q across tissues. Asterisks: Statistically significant. b: Schematic showing how tRNA-Q expands the codon decoding of tRNA-Asn. c: The design of the AAV vector carrying the codon mutated EGFP. d: Western blot of WT and mutant EGFP for each of the tRNA-Q decoded amino acids in the brain with quantification (densitometry). e: Western blot of WT and mutant EGFP for each of the tRNA-Q decoded amino acids in the liver with quantification. Asterisks indicate statistical significance. N = 5 animals per group.

Finally, we examined the ribosomal A-site pausing, which gives an indicator of the speed of decoding and codon optimality [Figure 6]. We compared the brain to each of the other tissues to elucidate the relative A-site pausing. Q-codons, except for Asp-GAC, were readily decoded in the brain compared to most other tissues indicating increased optimality and decoding speed [Figure 6a and b]. mcm5U codons showed variable optimality, with NAA mcm5U codons being readily decoded, while Gly-GGA and Ala-GCA having variable optimality or being suboptimal altogether [Figure 6a and c]. The results from A-site pausing were mostly in agreement with isoacceptors codon frequencies analysis, indicating that synonymous codon usage and bias is a determinant of decoding efficiency and mRNA translation across tissues.

### ⮚ tRNA modifications inform codon optimization algorithms for gene therapy

Given the clear links between tRNA modifications levels, tissue specific mRNA translation patterns, and codon usage and optimality, we asked whether the knowledge of tRNA modifications levels can inform codon optimization algorithms for optimal protein expression after gene or mRNA therapy. We focused on tRNA-Q modifications, given the clear enrichment of tRNA-Q in the brain vs other tissues [Figure 7a]. tRNA-Q expands the codon decoding from NAC codons to NAC and NAU codons (NAY codons) [Figure 7b]. Based on the data from our analysis, we focused on the brain and liver and example tissues to test the hypothesis. We argued that NAC codons would be decoded efficiently in either tissue, due to the existence of cognate anticodons. However, NAU codons will be less optimal in the Liver due to lower tRNA-Q levels. To test our hypothesis, we generated adenoviral vectors (AAV vectors) carrying wild-type (WT) or mutant EGFP [Figure 7c]. EGFP codon sequences were mutated from the original optimal NAC codons to NAU codons for each of the Q-modified amino acids tRNAs, or all four together [Supplementary table 3]. We used tdTomato as internal control. AAV vectors were injected intravenously into mice, and tissue samples collected after 3 weeks for western blotting validation of protein levels. In the brain, we did not observe changes in EGFP protein expression after mutating the codons [Figure 7d]. However, in the liver, we observed statistically significant downregulation of EGFP protein expression after mutating and of the Q-modified codons from NAC to NAU. This observation provides an indicator that a knowledge of tRNA modification levels could serve as an input for codon optimization strategies and can be further used for improving gene therapy efficiency.

## Discussion

In this work we present a comprehensive analysis of the tRNA epitranscriptome and identify its role in regulating codon usage and optimality across tissues. Importantly, our work highlights the important correlation between tRNA modifications and tissue specific codon decoding, signifying a role in fine-tuning mRNA translation and potential tissue specific physiologic roles. While several previous works have attempted to tackle this subject from various angles(26–28), our work represents the first comprehensive analysis of the various layers of the tRNA epitranscriptome and their direct correlation to translatomes to the best of our knowledge. We further provide proof of concept (PoC) data showing that knowledge of tRNA modifications in tissue, cell, or cell state specific context could be used to optimize gene therapies by fine-tuning the protein production *in vivo*.

At the tRNA modification levels, previous works by Guo et al(26) showed tissue specific enrichment levels but did not include the analysis of many anticodon modifications that are of great importance in fine-tuning codon optimality in tissues such as mcm5U, queuosine, or mitochondrial tRNA modifications. Further, previous works did not directly correlate tRNA epitranscriptome with codon decoding and translation(26, 27) or used limited metrics for codon analysis(28). Here, we directly analyzed the correlation between tRNA epitranscriptome on mRNA translation and codon decoding using multiple metrics. Our findings indicate either a role of tRNA modifications in fine-tuning mRNA translation via codon decoding, or physiologic adaptations of the tissues to meet the needs for synthesizing specific mRNAs that are codon biased. We also observed that tRNA modifications impacted codon usage and bias on two metrics: isoacceptors frequencies, and ribosome A-site pausing, both of which reflect codon bias and optimality.

tRNA modifications have been recognized as important players in many diseases and conditions(9, 11, 37). However, the cell-specific physiological roles are not explored for most known modifications. In our work, we identified queuosine (tRNA-Q) to be preferentially enriched in the brain and drive NAY codon recognition. Queuosine modifications were shown to play important role in regulating cellular response to arsenite induced oxidative stress and mitochondrial stress and dysfunction(15, 23), as well as playing a role in hippocampal neurons function and cognition(17). Interestingly, despite the reported importance of queuosine in codon decoding *in vitro*(*18, 23*), Knockout mice were phenotypically normal apart from cognitive and memory dysfunction(17). In addition, such distinct phenotype indicates the importance of tRNA-Q in brain function, in line with previous works that showed that Queuine, the precursor for queuosine, is protective against neurodegenerative diseases(38) and our observation of queuosine enrichment in the brain. This neural-specific phenotype sheds light as well on tissue and cell specific physiological functions of tRNA modifications that are rarely addressed. In the case of queuosine, it appears that the enrichment in the brain is vital for proper neuronal functioning, evident by Cirzi et al work(17). Such observations highlight the importance of understanding cell and tissue specific dependencies on tRNA modifications for proper translation in understanding tissue development, physiology, and pathologies.

Another interesting observation was the near uniformity of tRNA isoacceptors (anticodon) expression across tissues, in agreement with previous works(27). While it was shown previously by Pinkard et al(27) that there is tissue specific isodecoder tRNA level differences, tRNA isoacceptors expression were very stable across tissues. This agreement between our work and Pinkard et al(27) work despite using different tRNA sequencing techniques validates their findings, as well as ours. Importantly, while tRNA isoacceptors were shown to dynamically change in pathologies such as oxidative stress(6) and cancer(7) to impact codon decoding, it appears that this role is more or less stable in physiology. This was further show in recent work by Gao et al(33), where they showed that tRNA transcripts (isodecoders) vary between cell lines, but the anticodon pools (isoacceptors) are stable.

Codon usage and optimality have been a subject of immense interest, given their role in regulating many molecular processes such as mRNA stability and decay(39, 40), ribosome decoding fidelity(41), and mRNA translation efficiency(5, 42). Previous works revealed a plasticity of codon optimality that adapts to cell states. For example, during oxidative stress, tRNA modifications and tRNA expression dynamically change to alter codon optimality and drive translation of antioxidant proteins(6, 15, 23). This also occurs during cell proliferation and differentiation(43, 44). These findings alluded to the potential of tissue-specific codon optimality to regulate mRNA translation and local proteostasis. Such idea was challenged previously in Drosophila tissues by Allen et al(45). In their work, Allen et al(45) used codon mutated probes to evaluate the expression of rare codons in various tissues. They found that the testis and brain can express rare codons compared to other tissues. Attempts to elucidate tissue-specific codon optimality in mammalian tissues have been limited, and mostly relied on published datasets(46) and public databases(47), thus the analysis was usually limited by the data available. For example, no formal correlation between tRNA pools or tRNA modifications was made to explain codon optimality patterns or specific phenomena observed at the codon level in previous published works(46, 47), especially that the reliance on tRNA genes does not reflect actual tRNA transcript expression(13). In addition, the reliance on codon adaptation index (CAI) or tRNA adaptation index (tAI) might also not be ideal in such contexts where tissue specific metrics are needed, given that these two metrics depend on genomic information and not actual mRNA or tRNA levels(48, 49). This work presented herein represents the first analysis of mammalian codon usage and optimality that relies on tissue specific translational levels to deduce codon optimality to the best of our knowledge. Previously, Benisty et al(47) showed that most abundant mRNAs are A/T-ending biased. We observed the same pattern of enrichment in most tissues except for the brain and heart. However, while they made the argument that this bias is driven by tRNA expression profiles, we disagree on this notion. Mainly because we observed differences in codon decoding across tissues that cannot be explained by the stable expression of tRNA isoacceptors that we, and others(27), observed. In addition, Benisty et al(47) retrieved their tRNA data from another publication(50) which utilized TCGA datasets to deduce tRNA expression. However, we cannot deem the methods used to generate small RNA-seq datasets that were reported in the TCGA database to be fully capable to capture full length tRNA transcripts, and in turn, biases which are known to occur due to tRNA modifications and structure, could have played a factor. Thus, while we cannot negate the role of tRNA transcripts completely in driving the global A/T-ending codon bias, that role cannot explain the tissue specific patterns we observed, that could be explained by tRNA modifications.

Many important works have shown the role of tRNA modifications in regulating codon optimality and the decoding fidelity of ribosomes(4, 5, 9, 18, 25), we here provide evidence of such regulation at the tissue level in mammals. Our work, supported by previous work from Pinkard et al(27) and Gao et al(33), also shows that actual tRNA transcript expression does not appear to play important roles in driving codon optimality, given the uniformity across tissues. In our work, we also show a strong correlation between tRNA modifications and their decoded codons at the level of isoacceptors frequencies and A-site pausing, while this correlation weakens when we analyzed total codon frequencies across mRNAs. This further indicates that tRNA modifications drive mRNA translation via synonymous codon usage and bias. Codon optimization for gene and mRNA therapeutics have become a hot topic of research given the recent advances in mRNA vaccines, especially during the COVID-19 era(51).

Codon optimization was largely used to promote the stability and translation of bacterial and viral transcripts used for vaccination in mammalian cells(51). In that sense, genome-based metrics, such as CAI (codon adaptation index)(49) or tAI (tRNA adaptation index)(48) could be appropriate. However, we used a very different approach. We hypothesized that a knowledge of the synonymous codon usage, driven by tRNA modifications, could allow the tuning of protein expression of delivered gene constructs in different tissues. Indeed, we show here that a knowledge of tRNA-Q levels in the liver and brain allowed just that. This opens the door for optimizing gene and mRNA therapeutics in ways that allow maximum protein production in target tissues and cells while reducing the side effects from aberrant protein expression in other non-target tissues. It is important to note that, in future efforts to achieve such control, that diseased tissues have different codon usage patterns than healthy tissues, evident by a wealth of literature(11, 23). Thus, proper characterization and understanding of the relationship between tRNA epitranscriptome and codon usage in diseases is of utmost importance.

In conclusion, we show that tRNA modifications expression levels in various tissues are fine-tuned to drive tissue specific codon decoding and mRNA translation to maintain physiologic proteostasis needed for tissue functioning. We also show that a knowledge of tRNA modification levels and codon optimality in specific contexts is essential for better gene and mRNA therapeutic development. It is important to note that this work was somewhat limited as we did not conduct transcript specific tRNA modifications analysis, thus it remains unclear whether a modification that occurs in multiple tRNAs is enriched in all these tRNAs in a given tissue or if there some degree of fine-tuning. We also did not interrogate the levels of tRNA aminoacylation, which are known to play important roles in dictating translation and codon decoding(52). Future works should attempt to replicate our findings using different methodologies to ensure that these results are not due to specific technical limitations. It is also important to further clarify the physiologic roles of tRNA modifications in representative primary cells and not in commercially available cell lines that might not reflect the true physiology of tissues/cells.

## Methods

### Animals and sample collection, RNA isolation

Eight-week-old male C57BL/6J mice purchased from Kumagai Shigeyasu Shoten (Sendai, Japan) were maintained on a 12-hour light-dark cycle. For LC-MS/MS analysis, four mice were perfused with ice-cold PBS, and tissues (brain, lung, heart, liver, spleen, kidney, and muscle) were extracted immediately after perfusion and flash-freeze until RNA extraction. Tissues were homogenized with QIAzol (QIAGEN, #79306) and isolate small RNA using Purelink miRNA Isolation Kit (Thermo Fisher, #K157001). For small RNA sequencing and Ribosome profiling, three mice were perfused using ice-cold phosphate-buffered saline with 100 µg/ml cycloheximide (Sigma, #01810). Tissues were extracted immediately after perfusion and flash-freeze until RNA extraction. Tissues were homogenized with lysis buffer (20 mM Tris-Cl pH 7.5, 150 mM NaCl, 5 mM MgCl_2_, 1 mM dithiothreitol (Sigma, #D9779), 100 µg/ml cycloheximide, and 1% Triton X-100) and centrifuge at 13000 x g at 4°C for 15 min. For small RNA sequencing, 100 µl of supernatants were transferred to fresh tubes, and then 900 µl of QIAzol was added. Subsequently, small RNA was isolated using Purelink miRNA Isolation Kit, according to the manufacturer’s protocol. For Ribosome profiling, 500 µl of supernatant was kept for the downstream procedure. All animal experiments were conducted according to the procedures approved by the animal care facility of Tohoku University and according to the ARRIVE (Animal Research: Reporting In Vivo Experiments) guidelines. Ethical board approval was acquired prior to the commencement of this project.

### Small RNA quality control

Nanodrop one was used to analyze RNA concentration and purity. RNA integrity was analyzed using Bioanalyzer 2100 and the Agilent bioanalyzer small RNA kit (Agilent, #5067-1548). Example bioanalyzer traces are provided in supplementary figure 2.

### Quantitative analysis of tRNA modifications by Mass spectrometry

Quantitative liquid chromatography tandem mass spectrometry (LC-MS/MS) analysis of RNA modifications was done as reported previously with modifications(53). Small RNA enriched RNA fractions, which contained around 90% mature tRNA, were digested for 6 hours at 37 °C using a digestion mixture of: MgCl_2_ 2.5 mM, Tris (pH 8) 5 mM, Coformycin 0.1 μg/ml (Sigma, # SML0508), Deferoxamine 0.1 mM (Sigma, # D9553), Butylated hydroxytoluene 0.1 mM (Sigma, # W218405), Benzonase 0.25 U/μl (Sigma, # E1014-25KU), Calf intestinal alkaline phosphatase (CIAP; Sigma, #P5521) 0.1 U/μl, and Phosphodiesterase I (PDE I; Sigma, # P3243) 0.003 U/μl. Samples and standards were injected into Waters BEH C18 column (50 × 2.1 mm, 1.7 µm; Waters, #186002350) coupled to an Agilent 1290 HPLC system and an Agilent 6495 triple-quad mass spectrometer. The LC system was conducted at 25 °C and a flow rate of 0.3 mL/min. Buffer A was composed of 0.02% formic acid (FA) in DDW. Buffer B was composed of 0.02% FA in 70% Acetonitrile. The buffer gradient is shown in supplementary table 4. The UPLC column was coupled to an Agilent 6495 triple quad mass spectrometer with an electrospray ionization source in positive mode with the following parameters: gas temperature, 200 °C; gas flow, 11 L/min; nebulizer, 20 psi; sheath gas temperature, 300 °C; sheath gas flow, 12 L/min; capillary voltage, 3000 V; nozzle voltage. Dynamic MRM was used to detect modifications using transitions and collision energies listed in supplementary table 5. Peak areas were normalized to the sum of UV signal of the canonical nucleotides (U, C, G, and A) and expressed as area ratio. Four biological replicates were used per group.

### Small RNA and tRNA sequencing

We used methods that were described previously(54, 55) with some modifications. Briefly, to remove 3’ conjugated amino acids, 500 ng RNA sample was deacylated in 0.1 M pH 9.0 Tris-HCl buffer at 37℃ for 1 hour. Deacylated RNA was treated with AlkB demethylase (rtStar tRNA-optimized First-Strand cDNA Synthesis Kit, ArrayStar, #AS-FS-004) at ambient temperature for 2 hours to reduce tRNA methylation. RNA was purified using Zymo Directzol RNA micro kit (Zymo reserch, #R2060) and end repaired with T4 Polynucleotide Kinase (T4 PNK; NEB, #M0201) at 37°C for 30 min. All treated RNA samples were used for library preparation using NEBNext Multiplex Small RNA Library Prep Set for Illumina (NEB #E7300/7580) according to the manufacturer protocol. Size selection of amplified cDNA library was performed using 6% TBE PAGE in a range of 140 to 210 bp (microRNA to mature tRNA). Library concentration and quality were assessed by Bioanalyzer 2100 using DNA 1000 chip (Agilent, #5067-1504). All the samples were pooled and sequenced by Illumina Hiseq-X ten instrument in a pair-end, 150 bp read. Three biological replicates were used per group.

### Small RNA sequencing data analysis

Small RNA libraries were first processed by tRAX software(56) to perform adaptor trimming and alignment to mouse tRNA reference (mm10 genome) as well as quality control. tRAX was used to analyze mature tRNA and anticodons expression.

### Ribosome profiling

Ribosome profiling was performed as we previously reported(57), which was based on another protocol with few modifications(58). Ribosome foot-printing was performed on samples collected as mentioned above by adding 1.25 U/μl RNase I (NEB, #M0243L) to 500 μl clarified lysate and incubating samples on a rotator mixer for 45 min at room temperature. TRIzol reagent was added, and RNA was extracted using miRNeasy Mini Kit (QIAGEN, #217004). Ribosome protected fragments (RPFs) were selected by isolating RNA fragments of 27-35 nucleotides (nt) using TBE-Urea gel. rRNA depletion was conducted using NEBNext rRNA depletion kit v2 (NEB, #E7400L) followed by end-repair using T4 PNK after purification of the rRNA depleted samples using Oligo Clean & Concentrator Kits (Zymo research, #D4060). The preparation of sequencing libraries for ribosome profiling was conducted via the NEBNext Multiplex Small RNA Library Prep kit for Illumina according to the manufacturer’s protocol (NEB, #E7300S). Pair-end sequencing reads of size 150 bp were produced for Ribo-seq on the Ilumina Hiseq X-ten system. Three biological replicates were prepared per group.

### Ribosome profiling data analysis

Quality control for Raw Fastq was performed using FastQC. Next, adapter trimming and collapsing the pair-end reads into one was done using Seqprep. Trimmomatic(59) was used to further clean low quality reads. Bowtie2(60) was used to align the reads to a reference of rRNA and tRNA genes (mm10) to remove contaminants. After that, reads were aligned to the genome (mm10 downloaded from UCSC), reads counted using FeatureCounts(61), and differential expression conducted using Limma(62). Omics playgroud(63) was used for Ribo-seq data visualization and clustering as well as read normalization. Of note, we also used Stringtie(64) to generate TPKM normalized read counts which were comparable to the log2CPM generated from Omics Playground, thus we performed downstream analysis using the Omics playground output.

### Codon analysis

Isoacceptors frequency and total codon frequency were analyzed as we previously reported(34, 35). The amino acid normalized codon frequency is calculated as:

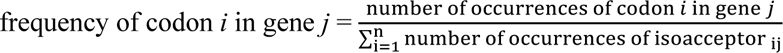

where isoacceptors refers to the synonymous codons for the amino acid encoded by codon *i* and n refers to the number of isoacceptors for a given amino acid (2 ≤ n ≤ 6).

The total codon frequency is calculated as:

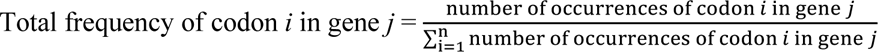

where n refers to the total number of codons used in the gene.

The t-statistic describing the codon frequency was calculated as:

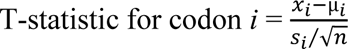

Where x_i_ refers to the mean frequency for codon *i* in the sample, µ_*i*_ refers to the mean frequency for codon *i* across the Mus musculus genome, s_i_ refers to the standard deviation of the frequency for codon *i* in the sample, and n refers to the sample size.

A-site pausing was calculated using Ribotoolkit(36) where the brain was compared to each tissue to deduce the relative ribosome pausing at each codon. Next, the output was converted into log2 fold change for data representation.

### Generation of adeno-associated viral (AAV) vector plasmids

The original EGFP (WT), and mutant EGFP with codons corresponding to each tRNA-Q modification (Tyr, His, Asn, Asp), or all four codons (All) in EGFP sequence modified from NA**U** to NA**C** were synthesized (Eurofins Genomics, Japan). To construct the vectors, CMV promotor and tdTomato sequences were obtained from pMuLE ENTR CMV tdTomato L5-L2(65) (Addgene #62152), and CMV promotor for EGFP was obtained from pMuLE ENTR CMV eGFP L3-L2(65) (Addgene #62143) by PCR amplification. All fragments were inserted into AAV backbone vector (pAAV-CAG-GFP, Addgene #37825) using NEB HiFi DNA Assembly Master Mix (NEB, #E2621), following the manufacturer’s protocol. The sequences for EGFP and mutated EGFP are presented in supplementary table 3. The following primers were used for CMV promoter and tdTomato cloning.

CTAGGAAGAGTACCATTGACGACATTGATTATTGACTAGTTATT

AATCAATGTCTAGAGGCTCGAGCTCTT

The following primers were used for CMV promoter cloning.

CGAGCCTCTAGACATTGATTATTGACTAGTTATT

GGTGGCTTTAGGATCCGA

The following primers were used for AAV backbone cloning.

TCGATATCAAGCTTATCGATAATCA

GTCAATGGTACTCTTCCTAG

### Viral vector production

Recombinant AAV vectors were produced through triple transfection of vector plasmids, pUCmini-iCAP-PHP.eB(66) (Addgene #103005), pAdDeltaF6 (Addgene #112867), into HEK293T cell, using 0.1% PEI MAX, pH 7.0 (Polysciences, Inc, #24765-100). Viral vectors were purified by ultracentrifuge as previously described(67). Briefly, cells were cultured with serum-free medium, and supernatant was collected 5 days after transfection. After virus concentration by PEG 8000 (Sigma, #P5413) and Benzonease treatment (Millipore, #70746-3CN), virus was purified by ultracentrifuge using Optiprep (Serumwerk Bernburg, #1893) discontinuous gradient. AAV titer was determined by qPCR method using AAVpro Titration Kit (for Real Time PCR) Ver.2 (Takara-bio, #6233) following the manufacturer’s protocol.

### *In vivo* virus transduction

6-week-old C57BL/6J male mice were injected intravenously with 1.0 × 10^11^ vector genome of AAV vector. Mice were sacrificed 3 weeks after transduction and tissues were extracted and flash frozen.

### Western blot

Tissue samples were collected and lysed with T-PER Tissue Protein Extraction Reagent (Thermo Scientific, #78510) containing cOmplete protease inhibitor cocktail (Roche, #4693116001). The tissue lysate was centrifuged at 16,000 x g for 20 min and supernatant was extracted for protein concentration measurement by Pierce BCA protein assay (ThermoFisher, # 23227). Each amount of protein was loaded into 8-16% Mini-PROTEIN TGX Precast Protein Gels (Bio-Rad, #4561106) and then transferred to Trans-Blot Turbo Mini 0.2 µm PVDF Transfer Packs (Bio-Rad, #1704156). The membranes were blocked by 5% skim milk (GFP; Wako, #198-10605) or PVDF Blocking Reagent for Can Get Signal (RFP; TOYOBO, #NYPBR01). Then, it was incubated overnight with primary antibodies GFP (Abcam, #ab6556), and RFP (MBL, #PM005) at 4℃. The band was detected with HRP-conjugated anti-rabbit antibody (CST, #7074S) and Pierce ECL Western Blotting Substrate (ThermoFisher, #32106). The protein bands were visualized by ChemiDoc MP (BioRad). The detected bands were analyzed using Image Lab Software (BioRad). One-way ANOVA with Dunnett’s multiple analysis and visualization was performed by GraphPad Prism 10 (GraphPad software).

### Data visualization and statistical analysis

Heatmaps and clustering and Pearson’s correlation analysis were conducted using Morpheus (https://software.broadinstitute.org/morpheus). Hierarchical clustering was performed using the “one minus Pearson correlation” metric and “average” as the linkage method. PLS-DA was conducted using the *R* language package mixOmics(68). Z-score analysis was conducted in *R*. ANOVA with Turkey’s post hoc analysis was conducted using SPSS v20 (IBM corp.).

## Data availability

Raw sequencing data were deposited in the sequence read archive (small RNA-seq: PRJNA1003133. Ribo-seq: PRJNA1004093). All other data are presented in the supplementary files. Further data can be provided by the corresponding author upon reasonable request.

## Supporting information

Suppmenetary tables

## Author contribution

**Ando D:** Sample collection and processing, sequencing libraries preparation, Animal experimentation, data analysis and interpretation, writing. **Rashad S:** Study conception and design, LC-MS analysis, Bioinformatics analysis, Data analysis and interpretation, writing, supervision, funding. **Begley TJ:** Codon algorithm development. **Endo H:** critically revised the manuscript. **Aoki M:** critically revised the manuscript. **Dedon PC:** LC-MS data interpretation. critically revised the manuscript. **Niizuma K:** critically revised the manuscript, Funding.

