## Supplementary figures for "tRNA modifications inform tissue specific mRNA translation and codon optimization"

**Supplementary tables legends:**

**Supplementary table 1:** Normalized peak areas of RNA modifications detected by LC-MS/MS. Related to figure 1.

**Supplementary table 2:** Differentially translated genes from the Ribo-seq dataset. Related to figure 3.

**Supplementary table 3:** Buffer gradient used for LC-MS/MS analysis.

**Supplementary table 4:** MRM table used for LC-MS/MS analysis.

**Supplementary table 5:** Sequences for EGFP and mutated EGFP used for AAV vector design

**Supplementary figure 1:** a: A compiled chromatogram showing all the detected modifications and their relative intensities. b: Variable (modifications) plotting from PLS-DA analysis. c: VIP (variable importance) plot from the PLS-DA analysis.

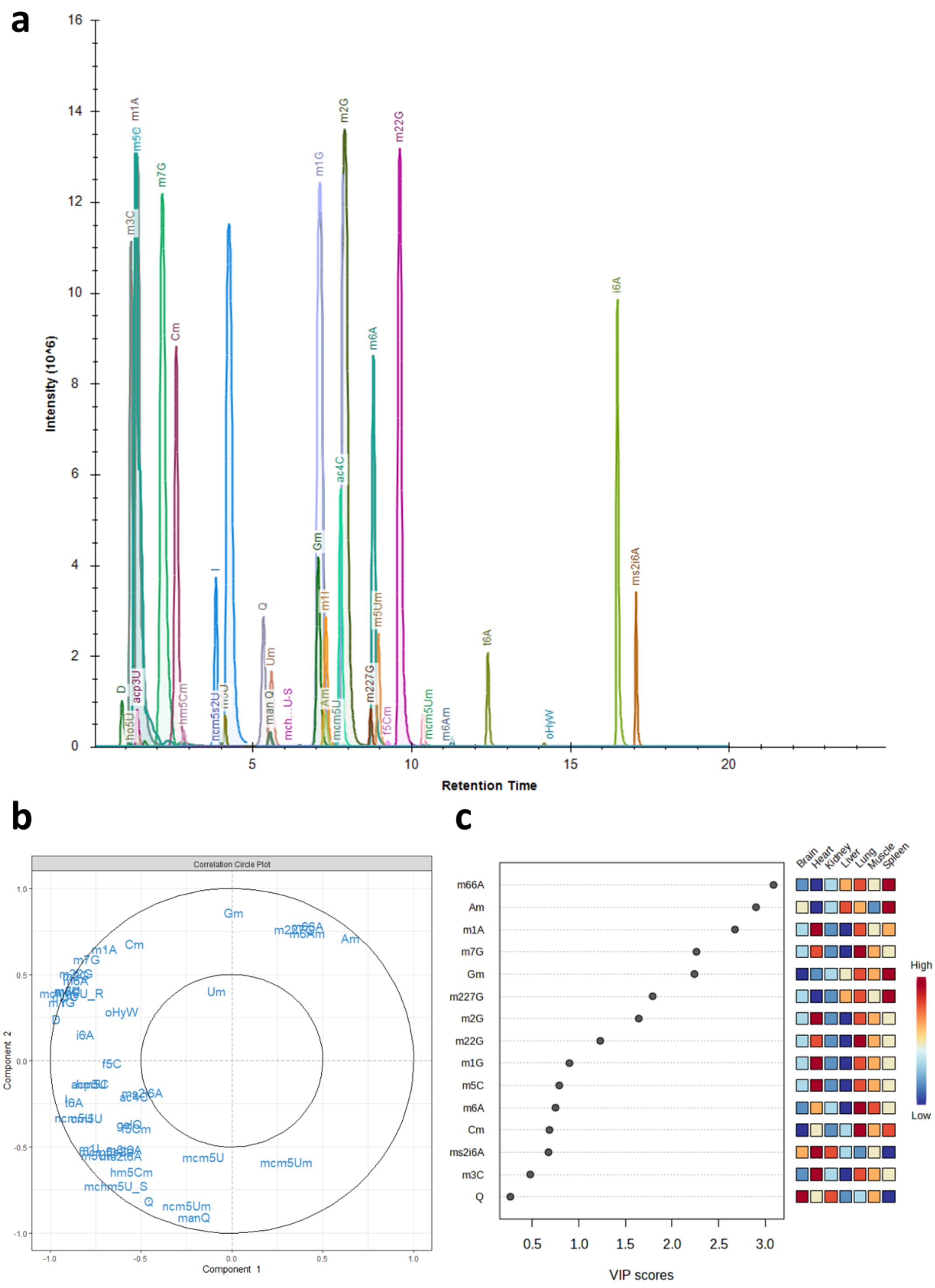

**Supplementary figure 2:** Example bioanalyzer traces showing the higher levels of non-coding RNAs in the spleen compared to tRNA levels.

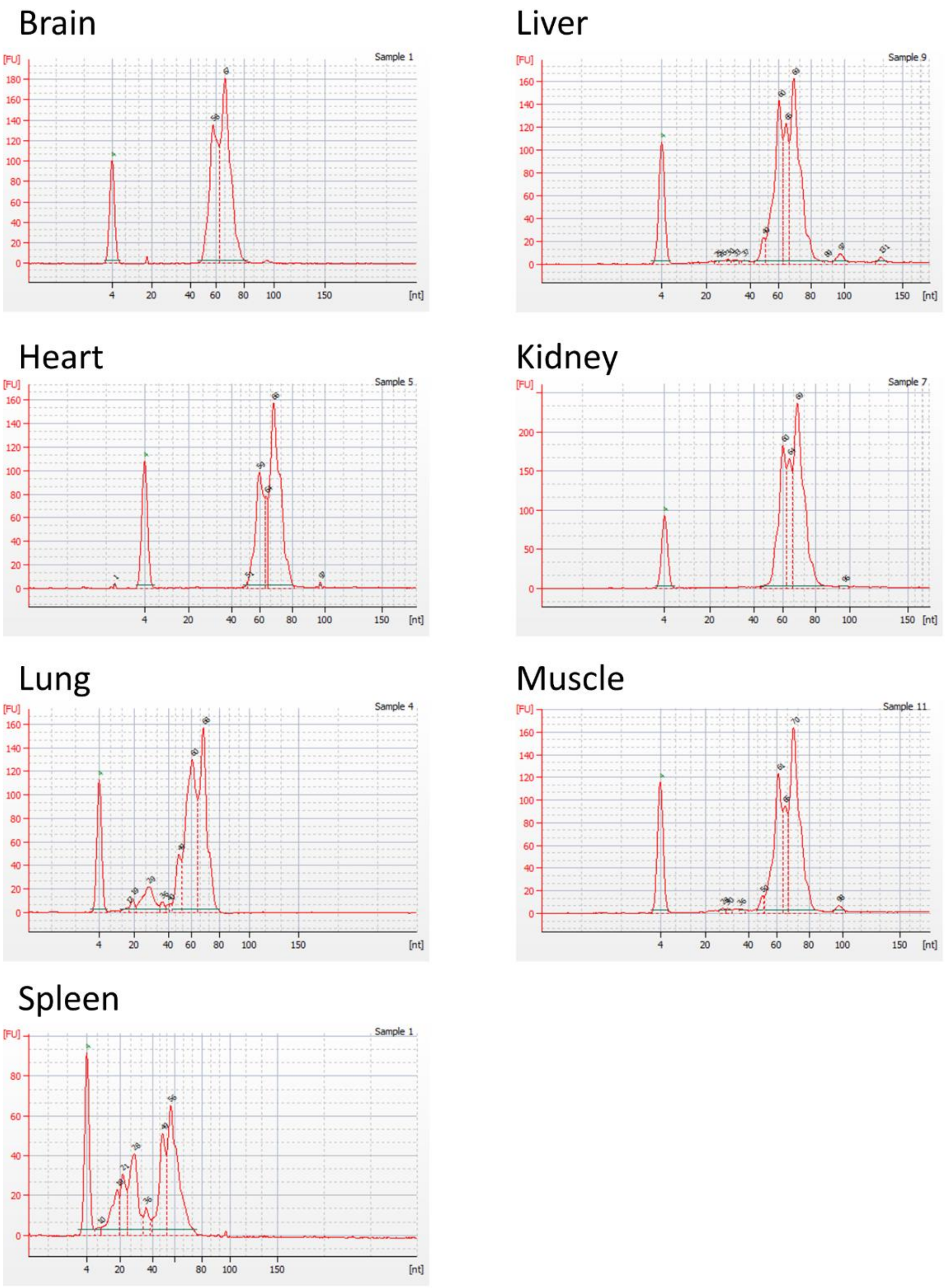

**Supplementary figure 3:** Analysis of mitochondrial tRNA expression. a: mt-tRNA expression by normalized read counts. b: mt-tRNA expression normalized by Z-score across tissues. c: Pearson's correlation analysis on mt-tRNA expression.

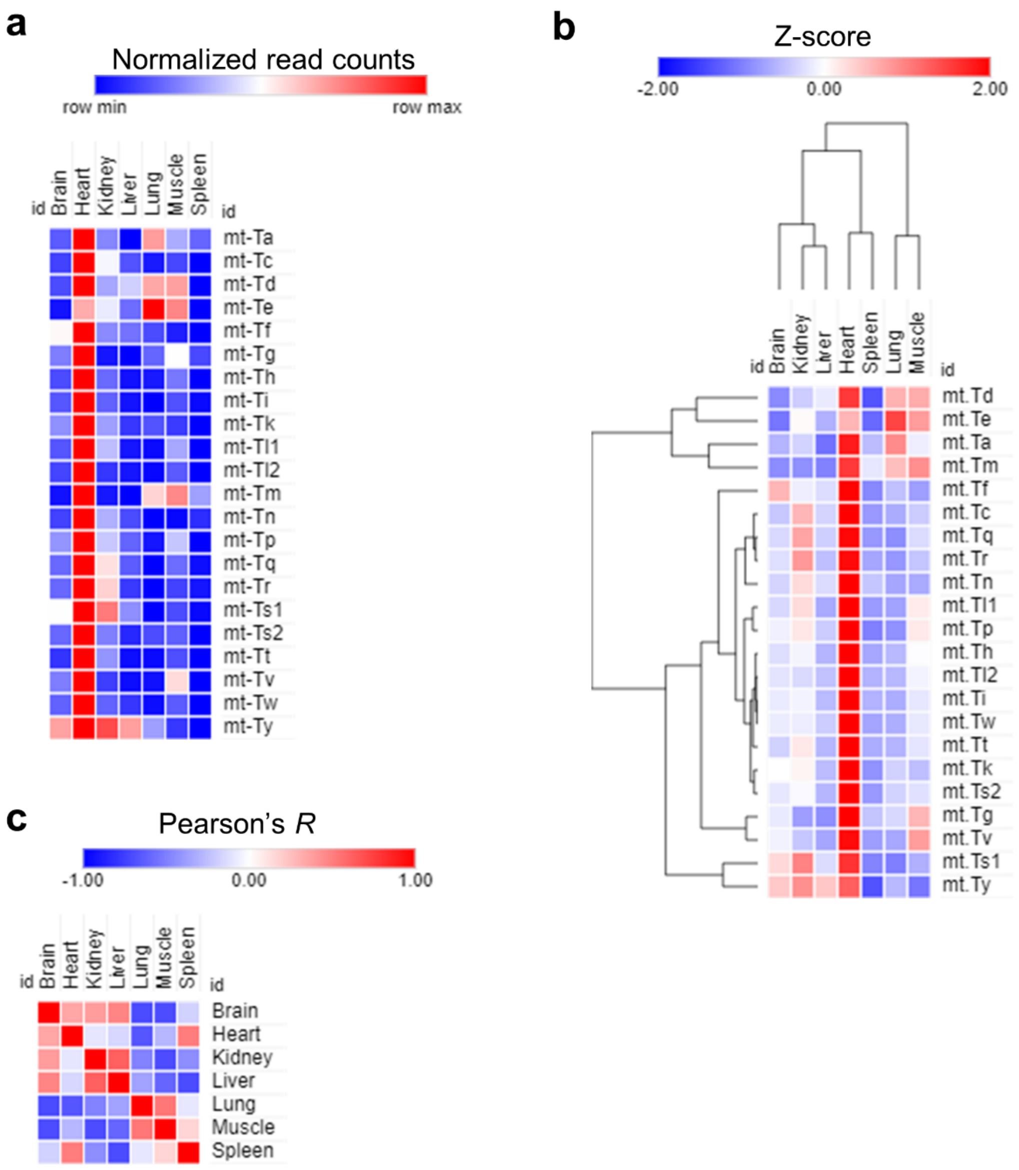

**Supplementary figure 4:** a: Annotations of heatmap clusters from figure 5a. b: GOBP UMAP clustering. c: Annotations of GOBP UMAP clustering.

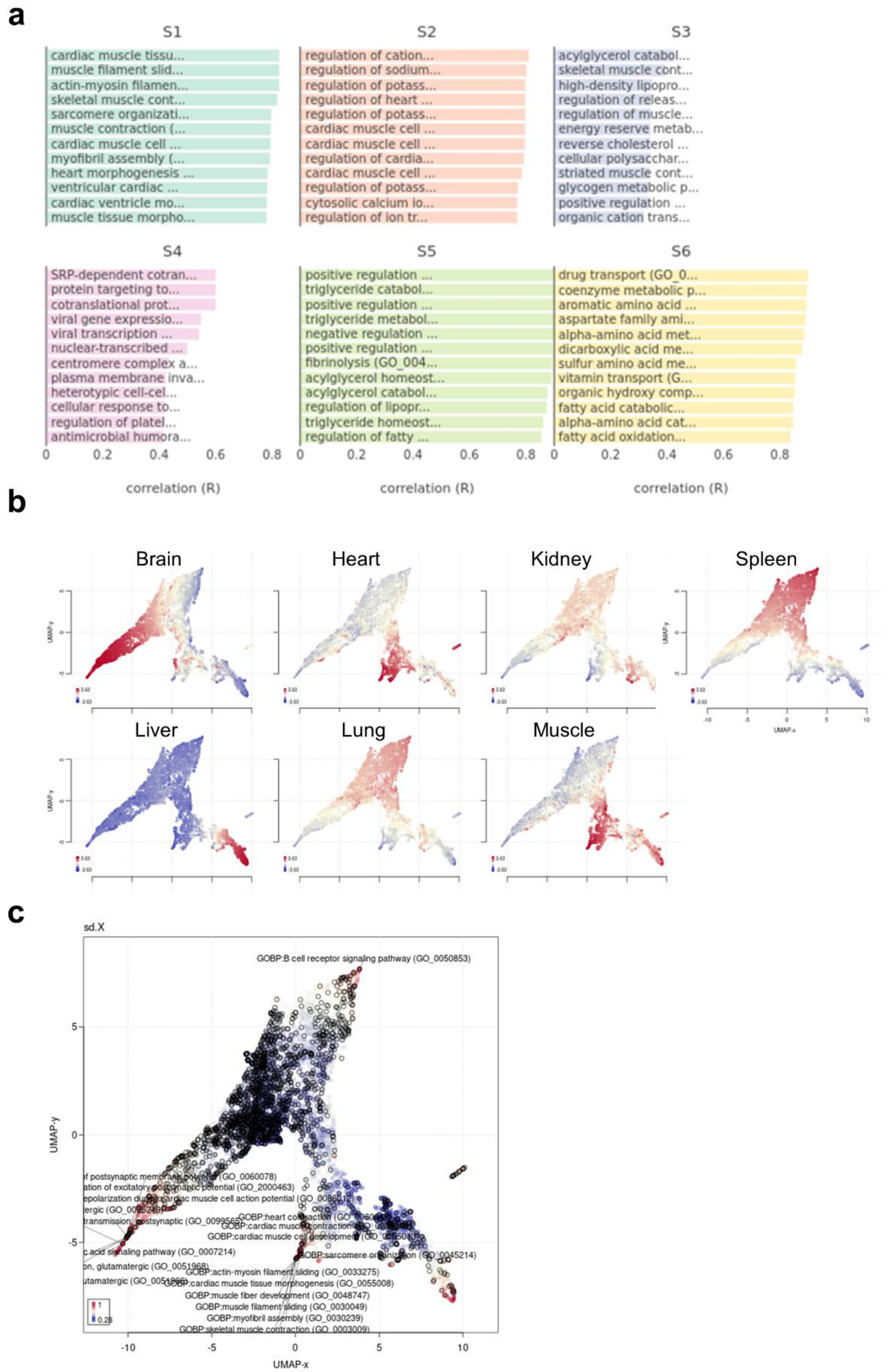

**Supplementary figure 5:** a: Phenotype (i.e., tissue type) based tSNE clustering. b: Phenotype annotation per sample. c: Phenotype annotation clustering patterns.

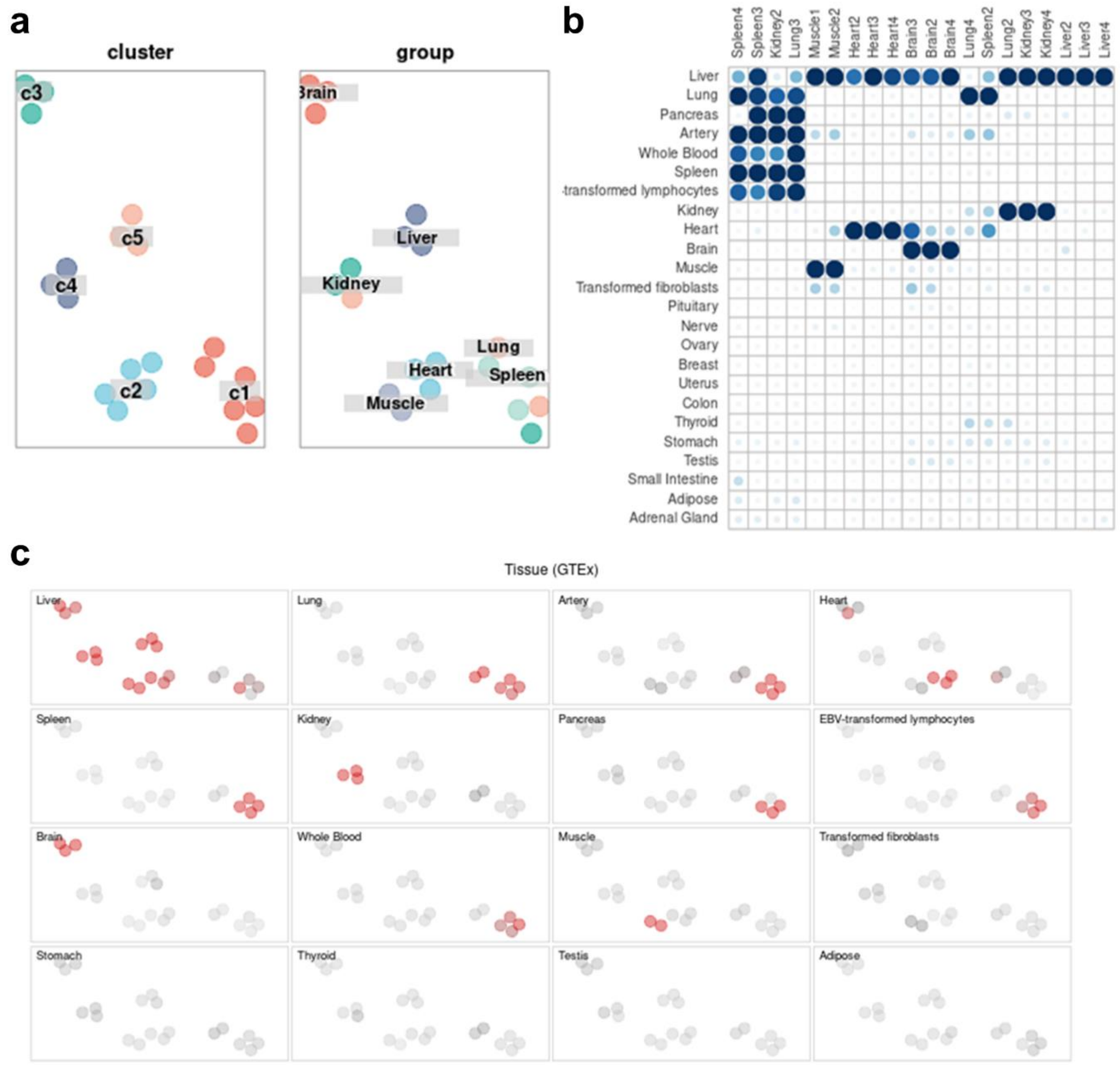

**Supplementary figure 6:** a: Translation pattern of RNA modifying enzymes in each tissue. b: PLS-DA analysis using the expression of RNA modifying enzymes as input.

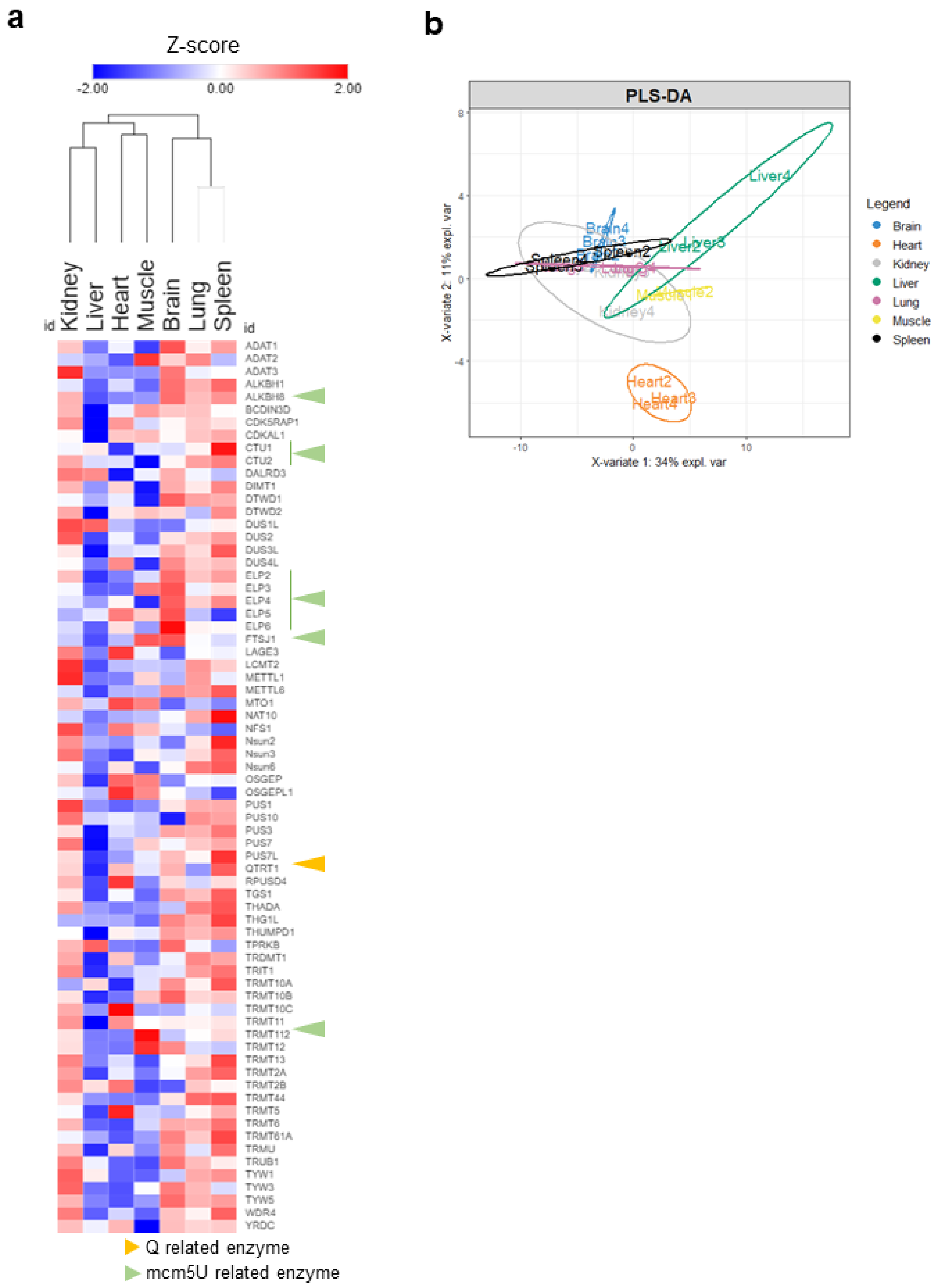

**Supplementary figure 7:** Different metrics for codon analysis used in this study.

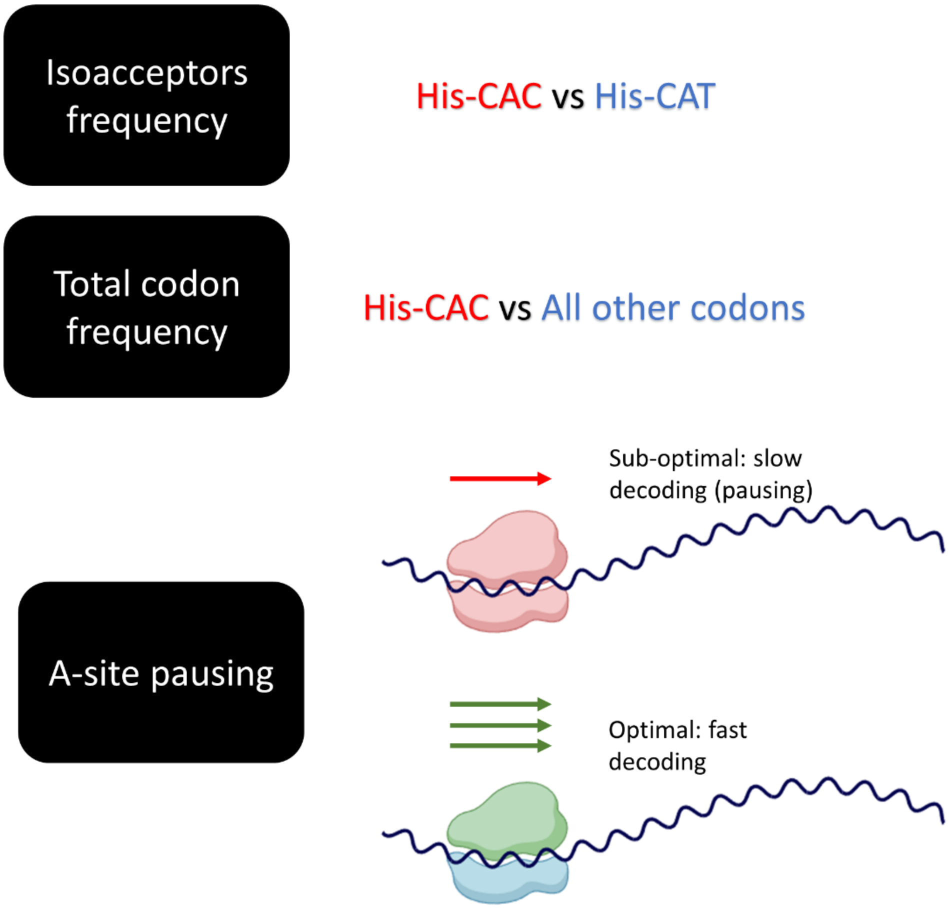

**Supplementary figure 8:** a: Pairwise correlation analysis (Pearson's correlation coefficient) of all codons from Figure 6a. b: PLS-DA plot of each codon using isoacceptors of the top 200 counted genes.

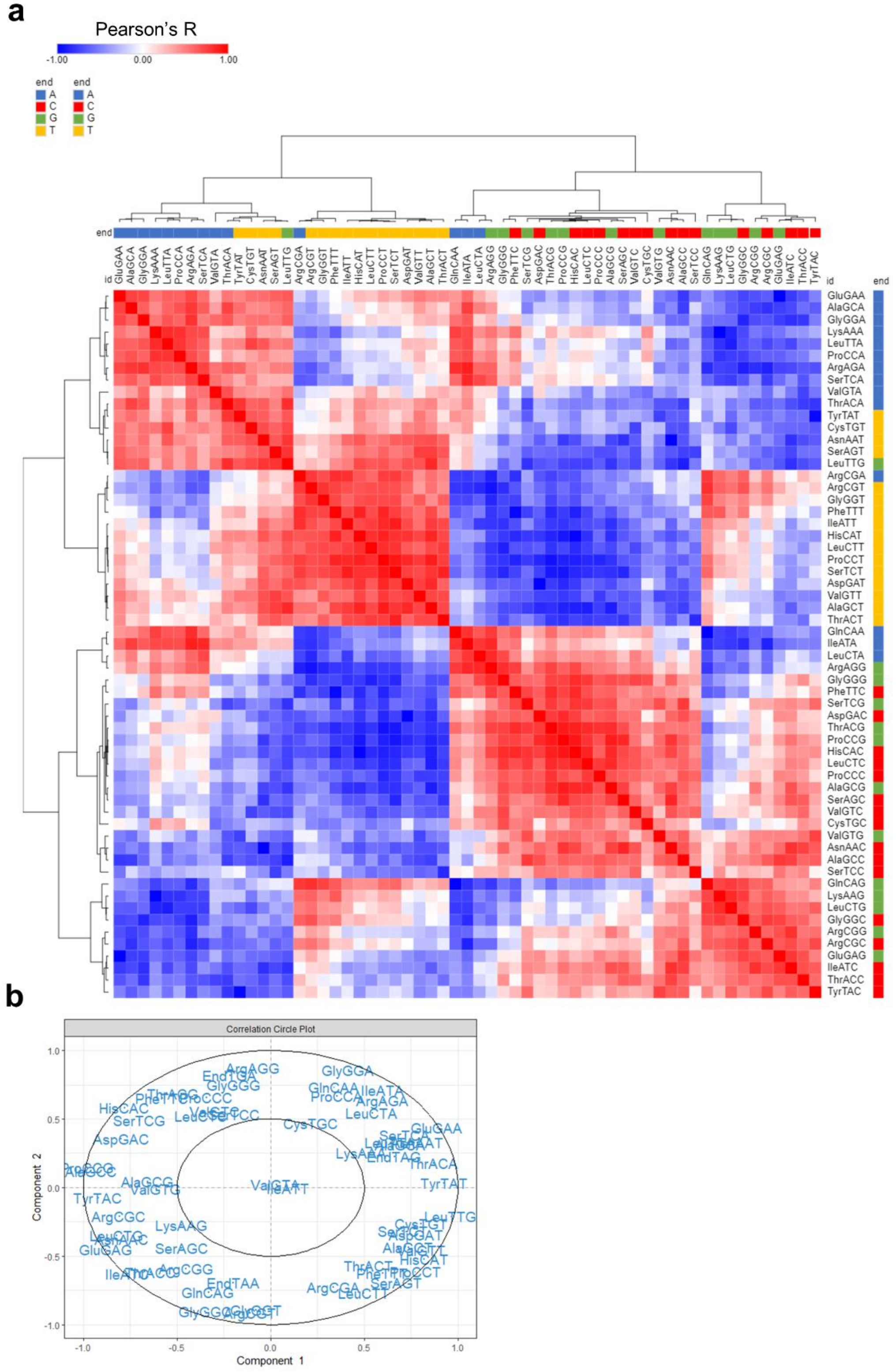

**Supplementary figure 9:** PLS-DA plot of each codon using total codons of the top 200 counted genes.

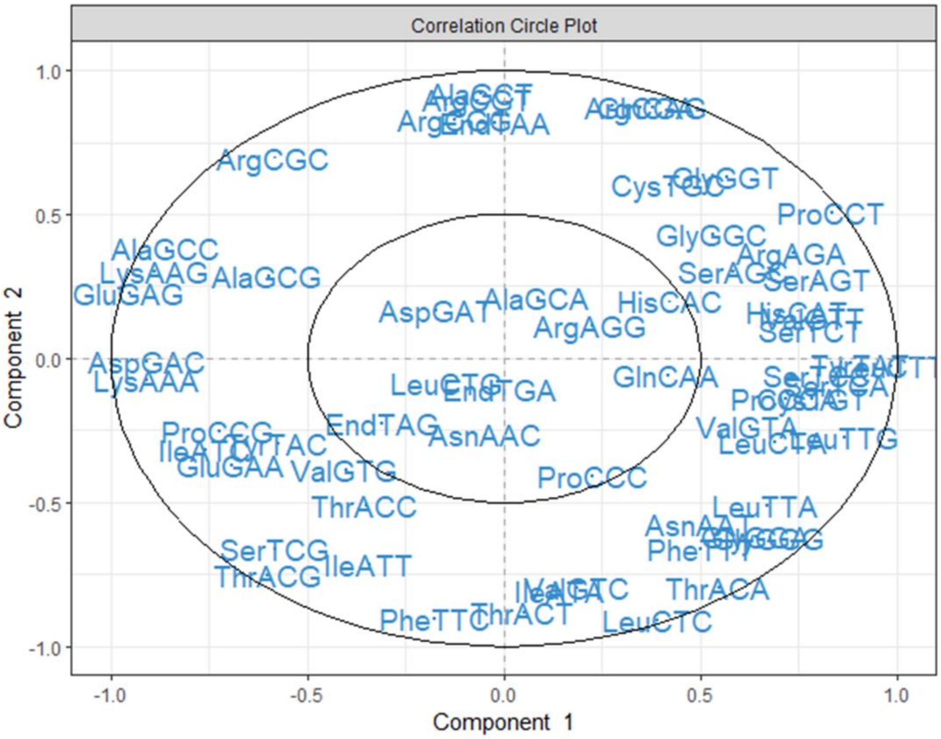
